# Single molecule dynamics of DNA receptor ComEA, membrane permease ComEC and taken up DNA in competent *Bacillus subtilis* cells

**DOI:** 10.1101/2020.09.29.319830

**Authors:** Marie Burghard-Schrod, Alexandra Kilb, Kai Krämer, Peter L. Graumann

## Abstract

In competent gram-negative and gram-positive bacteria, double stranded DNA is taken up through the outer cell membrane and/or the cell wall, and is bound by ComEA, which in *Bacillus subtilis* is a membrane protein. DNA is converted to single stranded DNA, and transported through the cell membrane via ComEC. We show that in *Bacillus subtilis*, the C-terminus of ComEC, thought to act as a nuclease, is not only important for DNA uptake, as judged from a loss of transformability, but also for the localization of ComEC to the cell pole and its mobility within the cell membrane. Using single molecule tracking, we show that only 13% of ComEC molecules are statically localised at the pole, while 87% move throughout the cell membrane. These experiments suggest that recruitment of ComEC to the cell pole is mediated by a diffusion/capture mechanism. Mutation of a conserved aspartate residue in the C-terminus, likely affecting metal binding, strongly impairs transformation efficiency, suggesting that this periplasmic domain of ComEC could indeed serve a catalytic function as nuclease. By tracking fluorescently labeled DNA, we show that taken up DNA has a similar mobility within the periplasm as ComEA, suggesting that most taken up molecules are bound to ComEA. We show that DNA can be highly mobile within the periplasm, indicating that this subcellular space can act as reservoir for taken up DNA, before its entry into the cytosol.

**Importance:** Bacteria can take up DNA from the environment and incorporate it into their chromosome in case similarity to the genome exists. This process of “natural competence” can result in the uptake of novel genetic information leading to horizontal gene transfer. We show that fluorescently labelled DNA moves within the periplasm of competent *Bacillus subtilis* cells with similar dynamics as DNA receptor ComEA, and thus takes a detour to get stored before uptake across the cell membrane into the cytosol by DNA permease ComEC. The latter assembles at a single cell pole, likely by a diffusion-capture mechanism, and requires its large C-terminus, including a conserved residue thought to confer nuclease function, for proper localization, function and mobility within the membrane.

## Introduction

*B. subtilis* is able to take up double-stranded DNA (dsDNA) from the environment at the onset of the stationary growth phase, e.g. under nutrient starvation conditions, and to integrate taken-up DNA into its chromosome. A multiprotein-complex is formed in 1-20% of the cells, dependent on the strain, and thus by heterogeneous expression within one culture, mediating the uptake of foreign DNA across the bacterial cell envelope (1–3). The competence-complex consists of the so-called late competence proteins, encoded by the *com*-operons *comE, comG, comF*, and *comC* (4–7). Expression of these operons depends on the transcription factor ComK (8), which is responsible for *B. subtilis* cells entering the so-called K-state, and is itself repressed by Rok (9). Rok is known to be a nucleoid associated protein, binding to A-T-rich regions, regulating a subset of genes associated with competence, but also other cell-surface and extracellular functions (10, 11). The expression of ComK increases in cells lacking *rok* from 10% to 60%, as demonstrated by GFP-fusions of ComK, leading to a much higher amount of cells entering the K-state (9).

Studies using epifluorescence-microscopy have shown that most of the major proteins essential for DNA-uptake localise to the cell poles in competent *B. subtilis* cells, such as ComGA, ComGB, ComEC, ComEB (12–14). It has been also shown that fluorescently stained dsDNA localises to the cell poles when incubated with competent *B. subtilis* cells. The labeled DNA can be integrated into the chromosome, leading to the formation of colonies displaying the particular resistance encoded by the transforming DNA (tDNA, (15)). These findings support the idea of a DNA-uptake complex located at the cell pole. Comparing data from other naturally competent bacteria, like the gram-negative *Vibrio cholerae* and the gram-positive *Streptococcus pneumoniae*, the concept of an uptake apparatus consisting of the competence complex and a type-II-/ type IV-secretion like pseudopilus has evolved in the field (16–18). In case of *B. subtilis*, its major component is thought to be the pseudopilin ComGC, facilitating the uptake of the tDNA through the cell wall (19).

The dsDNA-binding protein ComEA is an integral membrane protein that localizes throughout the membrane in a punctate manner (12, 14). It has been shown to be essential for binding of tDNA, carrying out the function of a DNA receptor during transformation (20, 21). An orthologue of ComEA, ComH of *Helicobacter pylori*, has recently been found to directly interact with the N-terminus of ComEC, the aqueous channel protein, which transfers the tDNA through the membrane into the cytosol (22, 23). ComH hands over the bound dsDNA to ComEC, while in case of *B. subtilis,* this role is taken over by ComEA. After being bound to ComEC, tDNA passes the channel as single-stranded DNA (ssDNA)(23), probably driven by a proton symport and the DEAD box helicase ComFA, providing energy by ATP hydrolysis for the process of transfer (6, 24, 25). Entering the cytosol, the exogenous DNA is coated by the ssDNA binding proteins DprA, SsbB and SsbA (26, 27), followed by integration into the chromosome through recombination proteins, RecA or RecN (13, 28, 29). How the ssDNA is generated inside the *B. subtilis* periplasm before entering the channel is still a matter of current investigations (18). In case of *Streptococcus pneumoniae*, the incoming dsDNA is hydrolysed by the membrane associated nuclease EndA (30), while for *Bacillus,* such a nuclease has not been discovered yet. One theory assumes that the late competence-protein ComEC takes over the aforementioned function (18).

ComEC is an integral membrane protein, possessing either 1 putative amphipathic helix, laterally associated with the membrane and 7 transmembrane helices according to Draskovic & Dubnau, (2005) (23), or possibly 11 transmembrane helices according to recent modelling studies of Baker *et al.* (2016). Draskovic & Dubnau (23) have investigated the membrane topology of ComEC by fusing different parts of the protein, either to LacZ or PhoA, determining the cytosolic or periplasmic localisation of protein domains based on the resulting enzyme-activity of the truncated LacZ-or PhoA-fusions. As a result, two soluble, periplasmic parts were identified. The N-Loop, located at the N-terminus of the protein, and the C-Loop, located at the C-terminus of ComEC. In addition, crosslinking-experiments of membrane protein fractions of competent *B. subtilis* cells were performed, identifying the formation of a dimer, probably by interaction of the two periplasmic parts (23). Recent *in silico* studies revealed the presence of a putative OB-fold, located at the N-terminus of the protein, which might be capable of ssDNA-binding, and a second domain at the C-terminus, putatively exhibiting exonuclease-function (31). These findings support the idea of ComEC providing the putative exonuclease among the competence-proteins, degrading the incoming DNA in order to pass the cell membrane. The ß-lactamase-nuclease-like domain of ComEC contains 2 conserved zinc-binding motifs, which are located within the periplasmic part of the protein, already described earlier by Draskovic *et al.* (2005). The model of Baker *et al.* (2016) (31) supports the existence of a large, soluble C-terminal part.

Earlier *in vivo* studies of ComEC report that in epifluorescence microscopy, the overall signal of a C-terminal YFP-fusion of ComEC is very weak or undetectable, indicating low abundance of the protein (12, 26, 32). Fluorescence-microscopy of two other competence proteins essential for DNA-uptake, ComGA and ComFA, revealed a similar localisation-pattern at the cell pole in competent *Bacillus* cells, combined with a non-uniform signal all over the cell. This prompted us to investigate the localisation of ComEC again by Epifluorescence-microscopy, but in addition by Single-molecule-tracking (SMT), in order to understand how its characteristic, polar localization pattern is achieved. To find out how the DNA uptake into the cell takes place, we truncated the C-terminal soluble part of the protein. In the absence of its soluble part, ComEC localises throughout the membrane, and the percentage of a population expressing ComEC increased. Removing its hydrophilic, C-terminal part slowed down the diffusion of the protein, indicating that ComEC assembles during competence of *B. subtilis* by diffusion/capture. In addition, analysing mutations of ComEC *in vivo* by determining transformation frequencies of several mutants revealed that the presence of a conserved aspartate residue, located within a putative zinc-binding motif of ComEC, is required for transformation. We have been able to determine the dynamics of taken up DNA at a single molecule level, and compared it to the mobility of ComEA, finding a similar diffusive behaviour. Our data suggest that exogenous DNA is directly bound by ComEA and can move within the periplasm before it is being transported across the membrane.

## Methods

### Growth conditions

*Escherichia coli (E. coli)* cells were grown in lysogeny broth (LB) medium and on LB-Agar-plates (1.5% Agar), supplemented with 100 μg/ml Ampicillin, at 37°C and 200 rpm. *B. subtilis* cells were grown on LB-Agar plates and in LB-medium at 30°C, supplemented with the required antibiotics, applying the following final concentrations: 5 μg/ml Chloramphenicol (Cm), 100 μg/ml Spectinomycin (Spec), 25 μg/ml Lincomycin and 1 μg/ml Erythromcyin (MLS), 25 μg/ml Kanamycin (Kan). In case an Amylase-assay had to be performed, LB-Agar was supplemented with 1% starch. If necessary, gene expression was induced by addition of either 0.05% Xylose (50% stock solution in ddH_2_0, sterile filtrated, used for C-terminal and N-terminal *mVenus*-fusions at the original locus) or 0.5 M IPTG (1 M stock solution in ddH_2_0, sterile filtrated, used for ectopic expression from the *amyE*-site), which was already included in the Agar-plates. For transformation of *B. subtilis*, a modified competence medium (MC-medium) was used after Spizizen, and cells were grown at 37°C, 200 rpm (33). A volume of 100 ml 10x MC-medium was composed as follows: 14.01 g K_2_HPO_4_ × 3 H_2_0, 5.24 g KH_2_PO_4_, 20 g Glucose, 10 ml trisodium citrate (300 mM), 1 ml ferric ammonium citrate (22 mg/ml), 1 g casein hydrolysate, 2 g potassium glutamate. Medium was sterile filtrated and aliquots of 1.5 ml were frozen at −20°C until use. Shortly before use, 1 ml of 10x MC-medium was added to a volume of 10 ml with 8.7 ml of sterile ddH_2_0 and 1 ml of 1 M MgCl_2_ (autoclaved) and supplemented with the corresponding antibiotics (or Xylose, if required). Frozen aliquots were not used longer than a week, as we found a loss of activity (transformation frequency) after long-term storage.

### Strain construction

All *B. subtilis* strains are listed and referenced in table S1. Strains of *E. coli* are listed in table S2 and corresponding Primer are listed in table S3. The C-terminal mVenus-fusion of ComEC (PG3817) was created by restriction-based cloning of the last 500 bp of the gene *comEC* into the vector pDL-mVenus via *Bgl*II and *Apa*I. pDL-mVenus encodes a MCS, followed by a 14 aa linker (GLSGLGGGGGSL) and *mVenus* (34). Inserting a *Bacillus* gene at the MCS of at least 500 bp, homology to the host genome is created, leading to a single-crossover of the vector with the *B. subtilis* chromosome. The gene of interest is thereby fused to the fluorophore at its C-terminus, still being located at its original locus. For C-terminal truncations of ComEC (PG3818, PG3819), the final protein sequence was chosen to be either 128 aa or 301 aa shorter than the full-length protein, removing either 1 putative helix and 1 zinc-binding-motif at position 667-672, aa: GDLEKE (PG3818), or two putative transmembrane helices and the C-terminal, soluble part of the protein, which is located in between the helices, and includes *both* putative conserved zinc-binding motifs at position 667-672 and 571-576, aa: GDLEKE and HADQDH (PG3819, (23, 31). Therefore, regions of 384 bp and 903 bp, starting from the C-terminus of *comEC*, were truncated from the template sequence for cloning. 500 bp of homology of the C-terminus were chosen, amplified and ligated to pDL_mVenus via *Eco*RI and *Apa*I restriction. After transformation of *B. subtilis* (PG001, see methods, transformation of *B. subtilis*), chromosomal integration was verified by PCR (see table S3, Primer EC fw and mV rev).

In order to investigate mutations of ComEC, the sequence of the gene was amplified and cloned via *Nhe*I and *Sph*I into pDR111 for ectopic expression at the *amyE*-site. In order to mutate amino acids H571, D573, D575, D668, and K671, plasmid DNA was isolated (GenElute Plasmid Miniprep Kit™, Sigma Aldrich) and further amplified by vector-PCR. Primer-pairs were designed back-to back with a size of 20 bp each, including the mutation at the beginning of the forward primer. through this technique, either one or two bp were exchanged, mutating the original amino acid to an alanine (used codons: *gcc, gca, gct*). Linearized plasmid DNA was phosphorylated and ligated (for detailed protocol see (35)), followed by transformation of *E. coli* DH5α. Plasmids were purified and sent for sequencing (Eurofins Genomics) to verify the presence of the desired mutations. In a final step, 24 bp encoding an RBS were added n-terminally to all constructs by a second vector-PCR (sequence, including **SD**: GATTAACTAATA**AGGAGG**ACAAAC (36, 37). Presence of the RBS was then verified by sequencing again. *B. subtilis* cells (PG001) were transformed by the resulting constructs. Ectopic integration at the *amyE*-site was proven by a 3x amylase-assay on starch-Agar. Therefore, plates were covered with 5 ml of Iodine-potassium iodide solution (Carl Roth) and incubated for 10 min at room temperature. Each time the assay was performed, a positive and a negative control were included. Finally, strains were transformed with chromosomal DNA of a PG3722, genotype: *ΔcomEC*. This resulted in strains encoding either the original gene or a mutated version of *comEC* at the *amyE*-site, and an erythromycin resistance cassette at the original locus instead of *comEC* (PG3917-PG3919, PG3927-PG3929). Presence of the resistance cassette was verified by PCR (see Primer Ery fw and Ery rev, table S3).

To create an N-terminal mVenus-fusion of ComEA (PG3909), 500 bp of the 5’ sequence of *comEA* were amplified and cloned into pHJDS-mV via *Apa*I and *Eco*RI, followed by transformation of *B. subtilis* PY79 (PG001). The original plasmid pHJDS1 (38) allows Xylose-inducible expression of N-terminal fusions of GFP at the original locus. In this case, *gfp* was exchanged for *mVenus* by restrictions-based cloning (PG3692, lab strain, unpublished, modified by Lisa Stuckenschneider via *Kpn*I and *Bam*HI). To facilitate SMT tracking of labeled DNA, we used a strain encoding a deletion of *rok* (PG876). In this background, *comEC* was deleted by transformation with chromosomal DNA of PG3722 to generate strain PG3816. Integration of the erythromycin resistance cassette was verified by PCR.

### Transformation of E. coli

To generate chemical competent *E. coli* DH5α cells for cloning procedures, a 200 ml culture was grown in LB medium at 37°C and 200 rpm, centrifuged at an OD_600_ of 0.6 and 3000 rpm, followed by resuspension in 15 ml of sterile filtrated, ice-cold 0.1 M CaCl_2_ solution, supplemented with 15% Glycerol. Aliquots of 150 μl were frozen in liquid nitrogen and stored at −80°C until use. For transformation, an aliquot was chilled on ice for approximately 5 min. 1 ng of plasmid DNA was added to cells, and incubated for 30 min on ice. After a heat shock for 90 sec at 42°C, aliquots were chilled on ice for another 10 min, followed by addition of 800 ml LB-medium and incubation at 37°C for 1 h (350 rpm). Cells were then plated on LB-Agar plates supplemented with the corresponding antibiotic and incubated at 37°C for 16 h.

### Transformation of B. subtilis & assays of transformation frequency

For transformation of *B. subtilis,* the application of glassware was obligate. Best results were obtained when strains were freshly streaked on LB-Agar, supplemented with inducing agents (0.05% Xylose, 0.5 M IPTG) and antibiotics, depended on the inserted plasmids or resistance cassettes, followed by incubation at 37°C over-night *before* transformation. From these LB-Agar plates, a 3 ml LB-over-night culture was inoculated and grown at 37°C and 200 rpm. 0.5 ml MC-medium, supplemented with antibiotics and required inducing agents (see Methods, growth conditions) was inoculated to an OD_600_ of 0.08, and grown at 200 rpm until the culture reached an OD_600_ of 1.5. Subsequently, 0.5-1 μg of chromosomal DNA, purified by isopropanol precipitation (35), was added to the cells and incubated further for 1-2 h at 37°C and 200 rpm. Cells were then plated on LB-Agar, containing the required antibiotics and incubated at 30°C for 48 h.

In order to perform assays of transformation frequency, 0.5 μg of chromosomal DNA were added to 0.5 ml of competent *B. subtilis* cells at an OD_600_ of 1.5 in a final concentration of 1 μg/ml. Further, cells were incubated for exactly 1 h at 37°C and 200 rpm. 100 μl of culture were diluted in 900 μl of LB-medium, followed by creation of a dilution-series in LB-medium. For calculating the number of viable cells/ml, a 10^−6^ dilution was plated on LB-Agar, while, depending on the strain, a 10^−1^, 10^−2^ or 10^−3^ dilution was plated in order to count colony forming units/ml (CFU). Finally, transformation frequency was calculated by dividing the CFU/ml by the number of viable cells/ml/μg DNA. For each strain, a technical triplicate and a biological duplicate was performed. Data were visualized via GraphPad Prism6 (GraphPad Software, San Diego, California, USA).

### Fluorescence staining of DNA

Preparation of labeled DNA was carried out following the exact protocol of Boonstra *et al.,* (2018). The modified nucleotide 5-[3-aminoallyl]-2’-deoxyuridine-5’-triphosphate (aminoallyl-dUTP, Thermo Scientific™) was incorporated by PCR using DreamTaq™ DNA Polymerase (Thermo Scientific™). A volume of 100 μl of dNTP-stock solution containing the modified nucleotide was set up as follows: 10 μl dATP (100 mM), 10 μl dGTP (100 mM), 10 μl aminoallyl-dUTP (50 mM), 10 μl dTTP (50 mM), 10μl dCTP (100 mM) 50 μl ddH_2_0. A 2300 bp fragment was amplified from the plasmid pDG1664 (PG311), using Primer prMB013 and prMB014 (15). The resulting PCR-product encoded for the erythromycin-resistance cassette of the vector including its flanking regions of the *thr*C-site, creating homology to the *B. subtilis* chromosome. PCR-reaction was carried out in a final volume of 400 μl. Subsequently residual template was removed from the PCR-reaction by a *Dpn*I digest for 1.5 h at 37°C. PCR product was purified (Qiagen PCR purification Kit) and eluted in 50 μl of 0.1 M NaHCO_3_ solution, pH 9. For staining of DNA, the fluorescent dye DyLight488 NHS Ester was used (Thermo Scientific™). In order to allow the aminoallyl-group of the incorporated nucleotide to react with the stain, facilitating the formation of an aminoallyl-ester, a 10x excess of dye was added to the PCR-product and incubated for 3 h in the absence of light at 25°C. Amount of dye was calculated as follows:

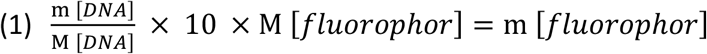

m= mass, M= molecular mass

Further, labeled DNA was purified from excess stain by PCR purification including an additional washing step (80% ethanol) and eluted in 50 μl ddH_2_0. In order verify staining and removal of residual dye, labeled PCR-product was detected in a 1% Agarose-gel by in-gel fluorescence at 488 nm (Typhoon TRIO, Amersham Biosciences, see figure S3). Aliquots were stored at −20°C under exclusion of light.

### Epifluorescence microscopy

*B. subtilis* cells were grown in MC medium to an OD_600_ of 1.5 at 37°C and 200 rpm (see growth conditions). 1% agarose pads were made from MC-medium. 100 μl of the melted agarose were sandwiched between two smaller coverslips (12 mm, Menzel) and let rest for 2 min. 3 μl of culture were dropped on a round coverslip (25 mm, Marienfeld), and covered with an agarose pad to fix the cells. A Zeiss Observer A1 microscope (Carl Zeiss) with an oil immersion objective (100 x magnification, 1.45 numerical aperture, alpha Plan-FLUAR; Carl Zeiss) was used for wide-field microscopy. Images were acquired by a charge-coupled-device (CCD) camera (CoolSNAP EZ; Photometrics) and an HXP 120 metal halide fluorescence illumination with intensity control. Fusions of ComEC and ComEA to mVenus were excited at a wavelength of 514 nm and detected at 727 nm. Cells were illuminated for 0.5-1.5 s at the mid-cell plane. Images were processed via ImageJ (39).

### Single-molecule tracking (SMT)

For single-molecule tracking, strains were grown under the appropriate conditions in order to obtain competent cells (see growth conditions). To study movement of mV-ComEA in the presence of DNA, 20 μg/ml chromosomal DNA (PY79) were added to cells entering stationary phase (t_1_) and incubated for 1 h, as described in Burghard-Schrod *et al.* (2020). In case labeled DNA was added for SMT, all steps were carried out in the dark. To analyse the diffusion of labeled DNA, 50 μl of competent cells were incubated with 20 μg/ml labeled DNA for 30 min on a shaking platform. Further, cells were centrifuged at low speed (500 rpm) and resuspended in 50 μl MC-medium (supernatant of the negative control, PG876). DNAse was added to the sample in a final concentration of 200 mg/ml and incubated for additional 40 min (15) at 37°C on a shaking platform. Cells were centrifuged and resuspended again in 30 μl of used competence medium. Slides for microscopy were prepared as described (see methods, Epifluorescence microscopy).

For SMT, an inverted fluorescence microscope (Olypmus IX71, Carl Zeiss Microscopy) was used. A 514 nm laser diode (100 mW, Omicron Laser) was used with 50% intensity, which enables slim-field microscopy of proteins fused to mVenus. To facilitate high-resolution detection of the signal of the fluorophore-fusions, an EMCCD camera (iXON Ultra EMCCD, Andor) was applied. Movies of 2500 frames were recorded by use of AndorSolis software in an exposure time of either 50 ms (PG3817-PG3819) or 30 ms (PG876, PG3816, PG3909). In case ComEC-mV was analysed at mid-cell plane, SMT was performed using a Nikon microscope equipped with an A = 1.49 objective and an Image-EMCCD camera (Hamamatsu) using an integrated autofocus of the microscope.

Bleaching curves of each movie were generated via ImageJ (39). To obtain signal only at the single-molecule level, intense signal of the fluorophore-fusions was removed by cutting the movies according to the bleaching curves, as strong signal could lead to a wrong connection of tracks, hindering further processing of the data. Typically, the first 500 frames of each movie had to be removed, to analyse only the frames covered by the horizontal part of the bleaching curve (indicating no constant bleaching of the signal was occurring anymore). Cell meshes were set with Oufti (40). Prepared movies were analysed with UTrack (41), while for each track a minimal step length of 5 was selected. Finally, tracks were analysed by SMTracker software (42). Diffusion coefficients and diffusive populations were calculated under use of the Gaussian-mixture model (GMM) and Squared displacement (SQD, for detailed description see supplemental methods).

## Results

### A C-loop deletion of ComEC localizes at the cell-membrane in epifluorescence microscopy

In order to study the influence of the C-terminal part on the function of ComEC, we created a C-terminal mVenus fusion of the full-length protein (PG3817) and to two versions truncated of the terminal 128 amino acids (AS), N648-N776, and secondly, of the whole C-terminal part predicted to be soluble, by removing the final 301 AS, L475 – N776. We named the fused truncations after the size of the truncated parts ComECΔ128-mV (PG3818) or ComECΔ301-mV (PG3819). Fusions were created at the original locus of the gene and were analysed by epifluorescence - microscopy of *B. subtilis* strain PY79 grown to competence. Initially, the location of these truncations was chosen by the presence of the transmembrane-helices described by Draskovic *et al.* (2005), depicted in figure 1, a). We expected a more soluble or fast-diffusing protein in case transmembrane-helices were removed, and/or no polar localisation pattern to be present anymore. The truncation ComECΔ128-mV displayed a polar signal similar to the wild-type fusion, but with fewer cells showing a signal at the cell pole, in 1.6 +/− 1.7% of the cells, compared to 5.7 +/− 2.05% (Fig. 1, b), Fig. 2) of the wild-type. Truncating a larger part of the C-terminus of ComEC in case of ComECΔ301-mV led to a punctate localisation pattern all along the membrane in 68 +/− 5.2% of the cells (Fig. 1, c), Fig. 2). The fact that the C-terminal truncation is restricted to the cell membrane strongly argues against a defect in membrane insertion caused by proteolysis. In case of ComEC-mV and ComECΔ128-mV, a large number of cells, 35.12 +/−1.15% and 47.67 +/− 1.2%, displayed a diffusive signal throughout the cells, which seemed to be reduced in the truncation ComECΔ301-mV, where the signal was either present at the membrane (and little in the whole cell), or not present at all (Fig. 2 a)). Still, in all three analysed strains, cells without signal were detected, in agreement with only a subpopulation of cells entering the state of competence. This can be seen in Fig. 2, where about 30% of the cells express ComK (as seen by CFP expressed from a ComK-regulated promoter), while 70% show no expression. Raw data and number of replicates are presented in table 1. Curiously, more cells than expected show a signal for localized or diffusive ComEC-mVenus (about 40%), which may be in a range expected for genetic noise, but almost 50% of the cells expressing the small ComEC truncation or more than 70% expressing the large truncation showed a fluorescence signal. While we have no explanation for this phenomenon, we note that a loss of the C-terminus of ComEC has an effect on the number of cells showing expression of the gene, or the entire *comE* operon.

**Table 1:** Localization of ComEC-mV and C-terminal truncations.

| Strain<br>genotype | PG681<br>*comK-CFP* | PG3817<br>*comEC-mV* | PG3818<br>*comECΔ128-mV* | PG3819<br>*comECΔ301-mV* |
| --- | --- | --- | --- | --- |
| number of cells (n) | 709<br>(224, 253, 232) | 307<br>(167, 140) | 475<br>(140, 151, 184) | 487<br>(158, 189, 140) |
| number of replicates (N) | 3 | 2 | 3 | 3 |
| % cell pole (+/-SD) | - | 5.74<br>+/-2.05 | 1.59<br>+/- 1.72 | 2.45<br>+/- 0.5 |
| % septum (+/-SD) | - | - | - | 4.67<br>+/- 0.54 |
| % diffusive (+/-SD) | 31.95<br>+/- 8.5 | 35.12<br>+/- 1.5 | 47.67<br>+/- 1.24 | - |
| % membrane (+/-SD) | - | - | - | 68.09<br>+/- 5.21 |
| % no signal (+/-SD) | 68.05 +/-<br>8.55 | 59.15 +/-<br>3.2 | 50.73<br>+/- 0.84 | 24.77<br>+/- 5.96 |

**Figure 1:**
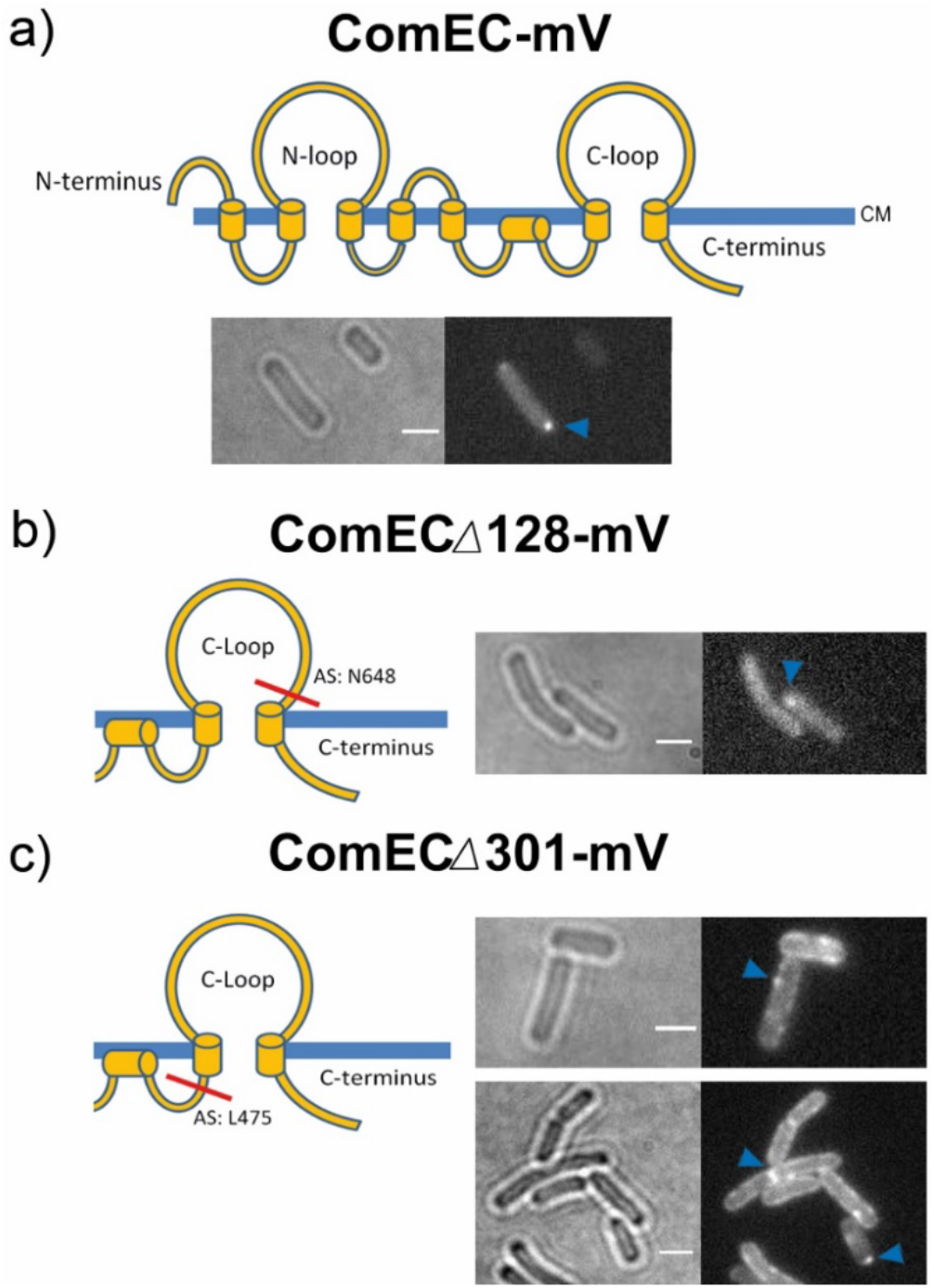
Epifluorescence-microscopy of ComEC-mV, ComECΔ128-mV, ComECΔ301-mV. Panel a) shows epifluorescence-microscopy of ComEC-mV (PG3817). The topology of ComEC is shown after Drascovic *et al.* (2005). CM indicates cell-membrane, the loops are thought to be present in the periplasm of the cell. In panel b) and c) the truncated versions ComECΔ128-mV (PG3818) and ComECΔ301-mV (PG3819) are shown. Cartoon indicate the position of each truncation at the C-terminus of ComEC (red line, b), c)). While the wild-type protein localizes at the pole (a)), a truncation of 128 AS leads to a more diffusive signal of the fusion and a rare signal at the pole (b)). A truncation of the whole C-loop of 301 amino acids localizes throughout the cell membrane (c)). Blue arrows indicate positions of high fluorescence. White bars 2 μm.

**Figure 2:**
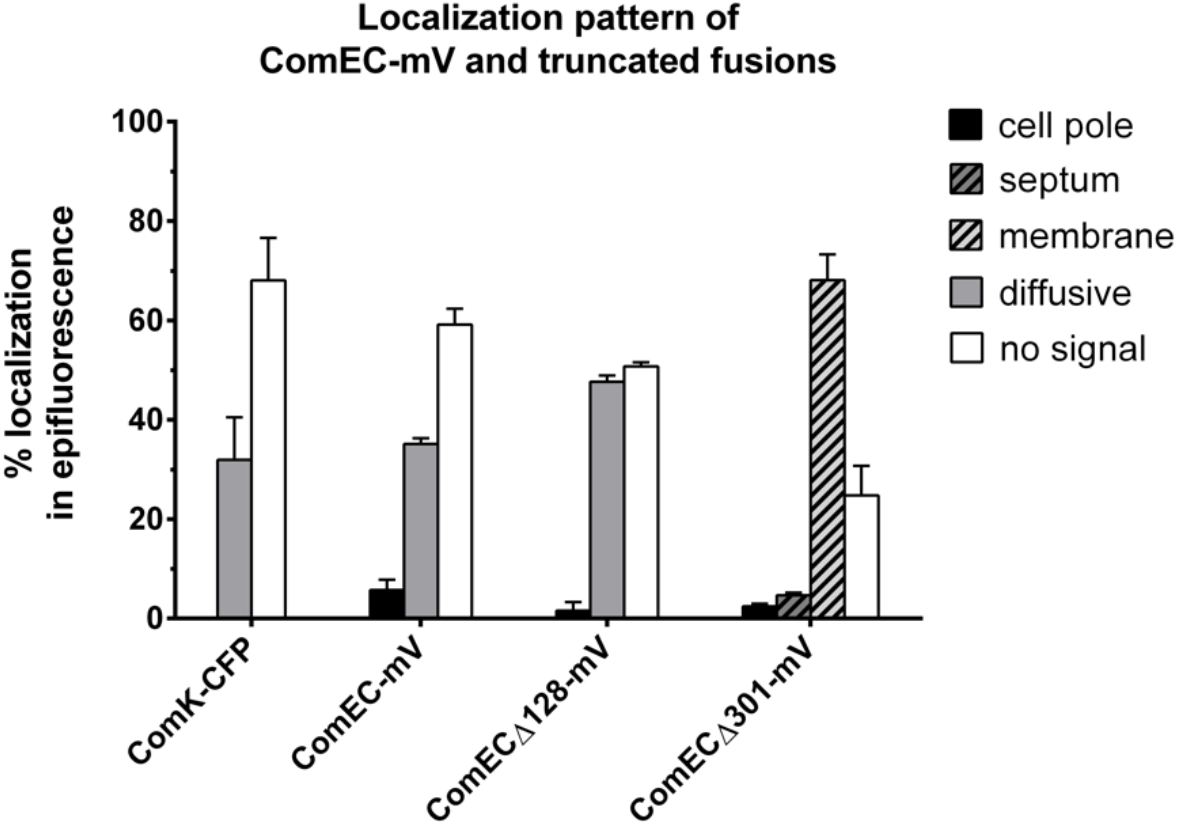
Localization pattern of ComEC-mV and C-terminal truncations. Different strains are indicated at the x-axis by their corresponding fluorophore-fusion (from left to right: PG681, PG3817, PG3818, PG3819). Bars represent the different cellular localization of mV-fusions as indicated by the y axis legend in %. In case of ComECΔ301-mV, strong septal or polar localization were counted separately, but always included a localization of the fusions at the cell membrane, and little diffusive signal throughout the cell. Error bars represent standard deviation of the mean. Number of replicates and raw data are listed in table 1.

### C-terminal truncations of ComEC lead to loss of transformability

Fusions were analysed with respect to their ability to take up DNA. Therefore, strains were transformed (see methods, transformation of *Bacillus subtilis* & transformation frequency) with chromosomal DNA of *B. subtilis*, encoding an Erythromycin resistance cassette (isolated from a Δ*comGA* strain, PG3717). Obtained colonies were selected for MLS-resistance, respectively. In case of a deletion of the whole C-loop of ComEC (PG3819, *comECΔ301-mV*) cells were hardly transformable (0.023% of the wild-type), while the mV-fusion of the full-length protein remained transformable, proving its functionality, as transformation frequency reached about 30% of the wild-type (see Fig. 3, table 2). Interestingly, activity was not completely abolished in the strain where we deleted only one helix and one putative zinc-binding site (PG3818, *comECΔ128-mV*), but reduced it about 1000 fold compared to the wild-type. Comparing the transformation frequency of ComECΔ128-mV (PG3818) to the full-length protein fused to mVenus, transformation frequency reached about 3% (see table 2).

**Table 2:** Transformation frequencies of strains expressing ComEC-mV and truncations.

| Strain<br>*genotype* | Transformation frequency<br>+/- SD | % wild-type<br>+/- SD | % ComEC-mV<br>+/- SD |
| --- | --- | --- | --- |
| PG001<br>PY79 | $2.93 \times 10^{-4} \pm 5.57 \times 10^{-5}$ | 100 | - |
| PG3817<br>*comEC-mV* | $9.32 \times 10^{-5} \pm 1.7 \times 10^{-5}$ | 31.78 +/-8.8 | 100 |
| PG3818<br>*comEC*<br>$\Delta 128\text{-mV}$ | $2.45 \times 10^{-7} \pm 9.3 \times 10^{-8}$ | 0.084 +/-0.031 | $2.63 \pm 9.9 \times 10^{-3}$ |
| PG3819<br>*comEC*<br>$\Delta 301\text{-mV}$ | $6.77 \times 10^{-9} \pm 1.05 \times 10^{-8}$ | 0.023 +/- $3.6 \times 10^{-3}$ | 0.072 +/- 0.01 |

**Figure 3:**
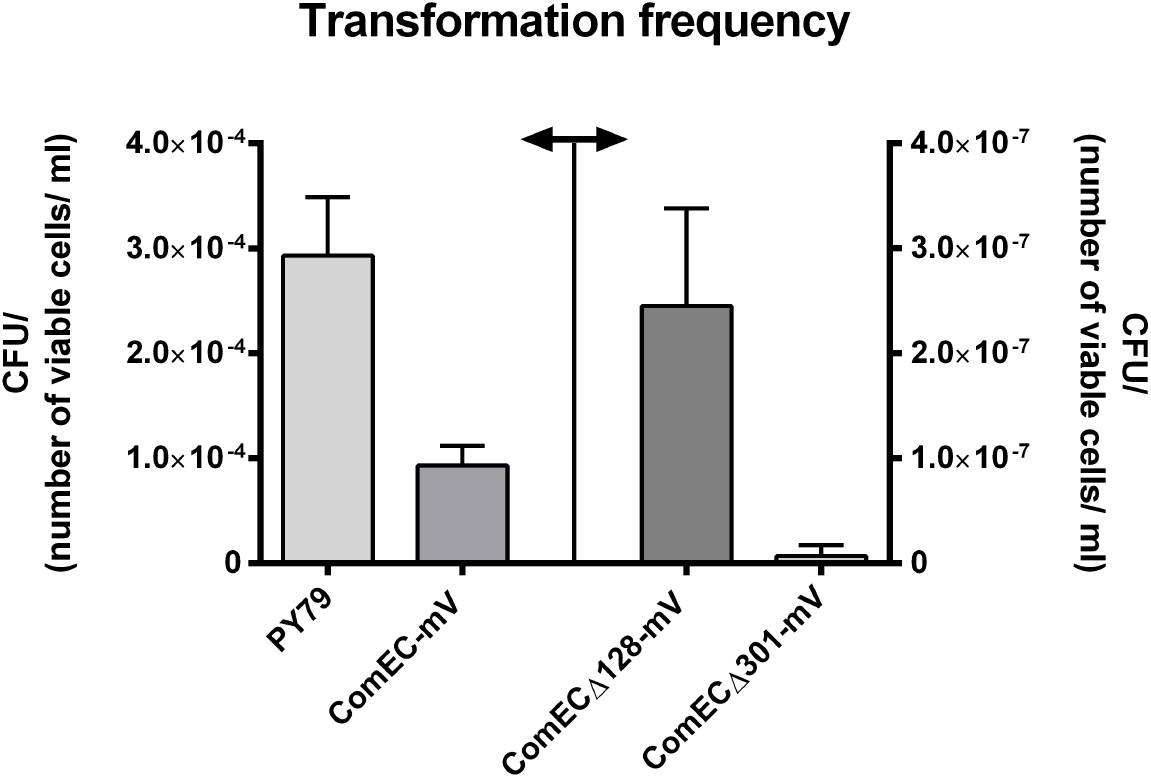
Transformation frequency of strains encoding *comEC-mV* and C-terminal truncations. Bar plot shows CFU divided by the number of viable cells/ml per μg chromosomal DNA/ml. Assays of transformation frequency were carried out for every strain (from left to right: PG001, PG3817, PG3818, PG3819) in biological duplicates (on two different days). Each measurement was performed in technical triplicates. Arrow indicates separation of the graph into two y-axes. Error bars represent the standard deviation of the mean.

### A conserved aspartate residue in the C-terminus of ComEC is essential for transformation

In order to further study the function of the C-terminal part of ComEC, we mutated conserved amino acids, which were postulated to be relevant for its putative exonuclease activity (Baker et al, 2016). Corresponding strains were analysed *in vivo* by a transformation frequency assay. Two conserved motifs identified by Baker *et al.* (2016) were (aa) **H**A**D**Q**DH** (motif 1, aa 571-576, conserved aa indicated in bold) and G**D**LEKE (motif 2, aa 667-671), which potentially coordinate two zinc ions and therefore could be directly relevant for functionality. We chose amino acids located in these motifs, and exchanged them for alanine (figure 4 a)). Genes were expressed ectopically from the *amyE* site, while the gene at the original locus was deleted. Mutating a histidine of the first motif (H571) lead to a very low transformability (12.73%), and mutation of a conserved aspartate of motif 1 (D573) abolished transformability completely (figure 4 b)). Exchanging the second aspartate of motif 1 (D575) resulted in a higher (but compared to the positive control, still low) transformation frequency of 30.5%. Mutating the aspartate located in motif 2 (D668), lead to a transformation frequency of 40% relative to the wild-type (PG3929), the highest value of all strains that were investigated, and where a putative zinc-binding amino acid has been exchanged. Interestingly, we found that transformation frequency decreased significantly in all mutations, except for one control, where we decided to mutate K671, which was postulated to be part of the conserved motif2 but *not* involved in zinc binding. In this case, transformation frequency reached 86% of the positive control (*ΔcomEC, amyE:comEC,* PG3929, see table 5 for transformation frequencies and percentages).

**Figure 4:**
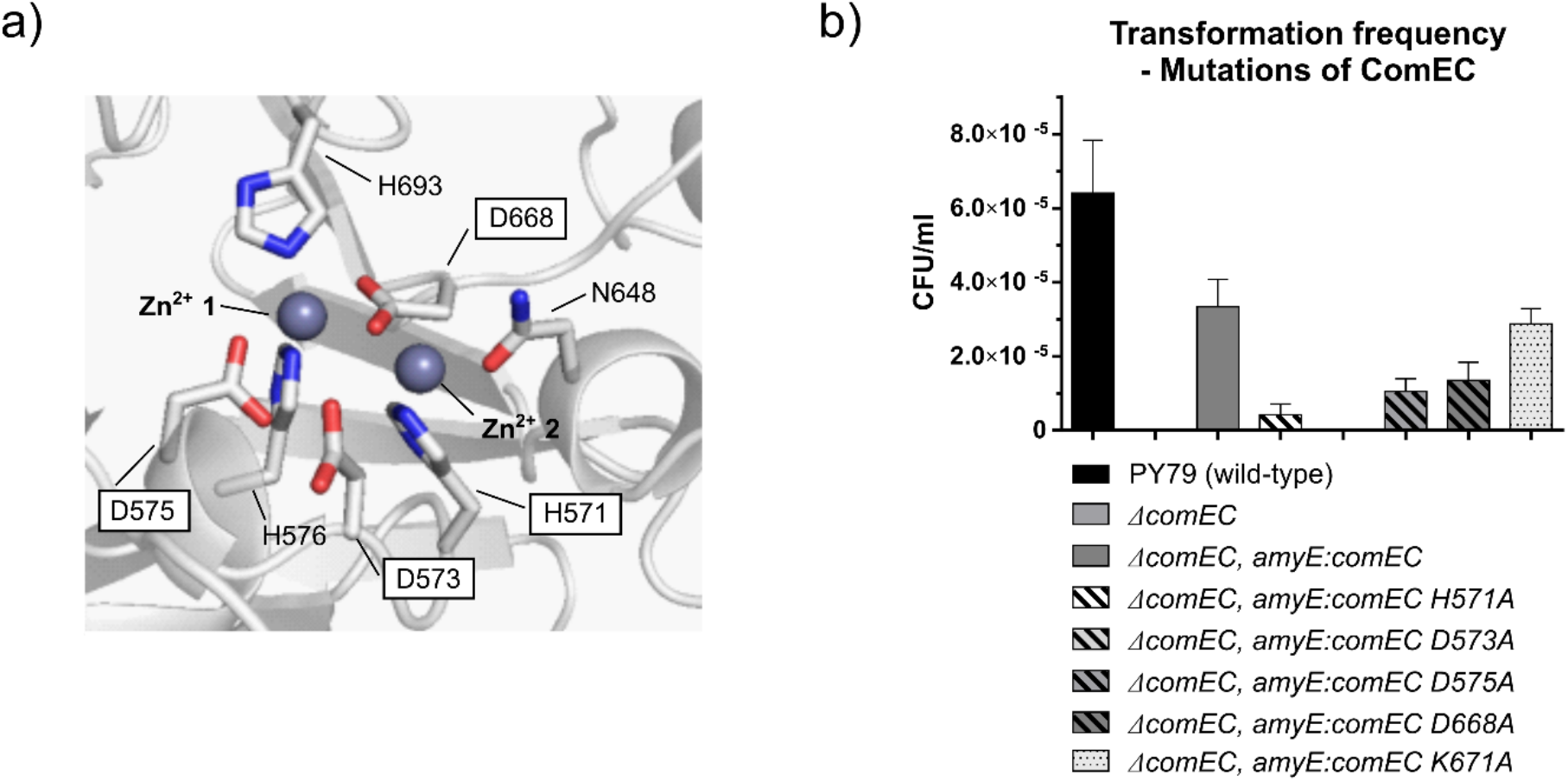
Transformation frequency of mutations of ComEC. In a) the active site of the β-lactamase domain of ComEC is shown, based on the model of Baker *et. al.,* (2016). According to the model, two zinc ions (Zn^2+^ 1, Zn^2+^ 2) are coordinated by two conserved aspartate residues (D575, D668), one arginine (N648) and three histidine residues (H571, H576, H693). The mutated amino acids are indicated by white boxes. Respective residues were exchanged by an alanine. Mutated genes were integrated and induced from the *amyE*-site, while the original gene had been deleted, in order to study the effect of the mutations on transformation. b) shows transformation frequency of the corresponding strains, while genotype is indicated by the legend. As additional control, amino acid K671, which is located close to the conserved residue D668, has been exchanged for an alanine. We found that the conserved residue D573, which appears to be close to the active site and present in the conserved motif **H**A**D**Q**DH**, but originally not depicted in the model of Baker *et al.* (2016), is essential for transformation. Assays were performed in technical triplicates and in biological duplicates. Error bars indicate standard deviation of the mean.

### *B. subtilis* ComEC moves within the membrane during the state of competence

In order to investigate the localisation of ComEC, we decided to analyse the fusion ComEC-mV by single-molecule tracking (SMT) in *B. subtilis* cells, grown to competence. Therefore, we applied an exposure time of 50 ms, where we made different observations by focusing on different cell planes during microscopy. By focusing to the central plane of the cell, we found a clear localisation pattern of ComEC-mV along the membrane (Fig. 5a), lower panel, movie S1), where we found mobile molecule tracks (shown in blue) or molecules moving within a confined area (red, most pronounced at the cell poles) at the cell periphery (green tracks show transitions between diffusive and constrained motion). However, by focusing on the upper level of the cell, we gathered more tracks, because molecules moved all along a bent surface (Fig. 5b). In both cases, clear large foci were visible, which for the upper level focus were found at all cellular places, since we were looking onto the surface of the cell. Accumulation of ComEC at the cell poles was not seen using the upper cell focus (figure 5b)), but diffusive dynamics of the protein in x and y direction was much better captured in this mode, rather than following movement in largely only x direction along the membrane, as in the central focussing mode. We therefore decided to determine velocity of our fusions at the upper-cell level. Applying Gaussian-mixture-model (GMM) analyses, we fitted 2 diffusive populations to the overall movement of the full-length fusion, which resulted in an overall well representation of the obtained step lengths. The two population fits showed that ComEC molecules fall into two modes of movement, one static and one mobile population, having diffusion-coefficients of 0.017 +/− 0.00013 μm^2^/s and 0.31 +/− 0.0011 μm^2^/s. The latter mobility corresponds to that of a membrane protein with several transmembrane spans (34). We found a large diffusive fraction in case of the full-length protein fusion, ComEC-mV, comprising 87.1% of all detected movements (see table 3). Comparing the sizes of the two diffusive fractions of the C-terminal truncations of ComEC, we found a decrease of the mobile fraction and an increase of the static fraction when the protein was truncated at its C-terminus in general. We found that ComECΔ128-mV showed a smaller mobile population of about 63% (24% more static than the wild-type), while for ComEC301-mV, we detected even fewer mobile tracks of a population of 49.8%. As already indicated by its appearance in epifluorescence (figure 1), we found a large difference comparing the full-length protein fused to mVenus and ComECΔ301-mV, a 37.3% decrease of the mobile fraction when the whole C-terminus was deleted. We infer that our fusions are stable proteins, because free mVenus (or any fluorescent protein) has not been reported to localize at the membrane on its own, and all our fusions show clear membrane localisation in SMT or epifluorescence. Unfortunately, we were unable to visualize ComEC-mVenus or mutant proteins by Western Blotting. Thus, we cannot exclude that we have been visualizing proteins lacking a part of their N-terminus, although this is unlikely because ComEC-mVenus is a functional protein fusion. We did not detect tracks in all cells as fusions were only expressed in cells being in the K-state, and therefore calculated the number of tracks per cell on the percentage of cells where tracks were present. Here we found no difference in case of ComEC-mV compared to ComECΔ128-mV (9.84 tracks, see table 3), but a moderate increase of tracks per cell in case of ComECΔ301-mV (11.8 tracks). These results indicate an increase of signal at the membrane in general when a part of the C-terminus was deleted (see overlay of tracks on cell, suppl. Fig. 1, in agreement with the epifluorescence analyses).

**Table 3:** Single-molecule tracking of ComEC-mV, ComECΔ128-mV and ComECΔ301-mV.

| Strain | PG3817<br>*genotype:*<br>*comEC-mV* | PG3818<br>*genotype:*<br>*comECΔ128-mV* | PG3819<br>*genotype:*<br>*comECΔ301-mV* |
| --- | --- | --- | --- |
| D<sub>1</sub> +/- SD [$\mu\text{m}^2/\text{s}$]<br>(static population) | 0.017 +/- 0.00013 | | |
| D<sub>2</sub> +/- SD [$\mu\text{m}^2/\text{s}$]<br>(mobile population) | 0.31 +/- 0.0011 | | |
| Pop<sub>1</sub> +/- SD [%]<br>(static population) | 12.9 +/- 0.14 | 37.5 +/- 6.64 | 50.2 +/- 4.38 |
| Pop<sub>2</sub> +/- SD [%]<br>(mobile population) | 87.1 +/- 0.14 | 62.5 +/- 6.64 | 49.8 +/- 4.38 |
| Analysed cells | 165 | 106 | 153 |
| Tracks (total) | 1280 | 915 | 1617 |
| Cells with tracks | 130 | 93 | 137 |
| Cells with tracks (%) | 78.78 | 87.73 | 89.54 |
| Tracks/cell | 9.84 | 9.84 | 11.8 |

**Figure 5:**
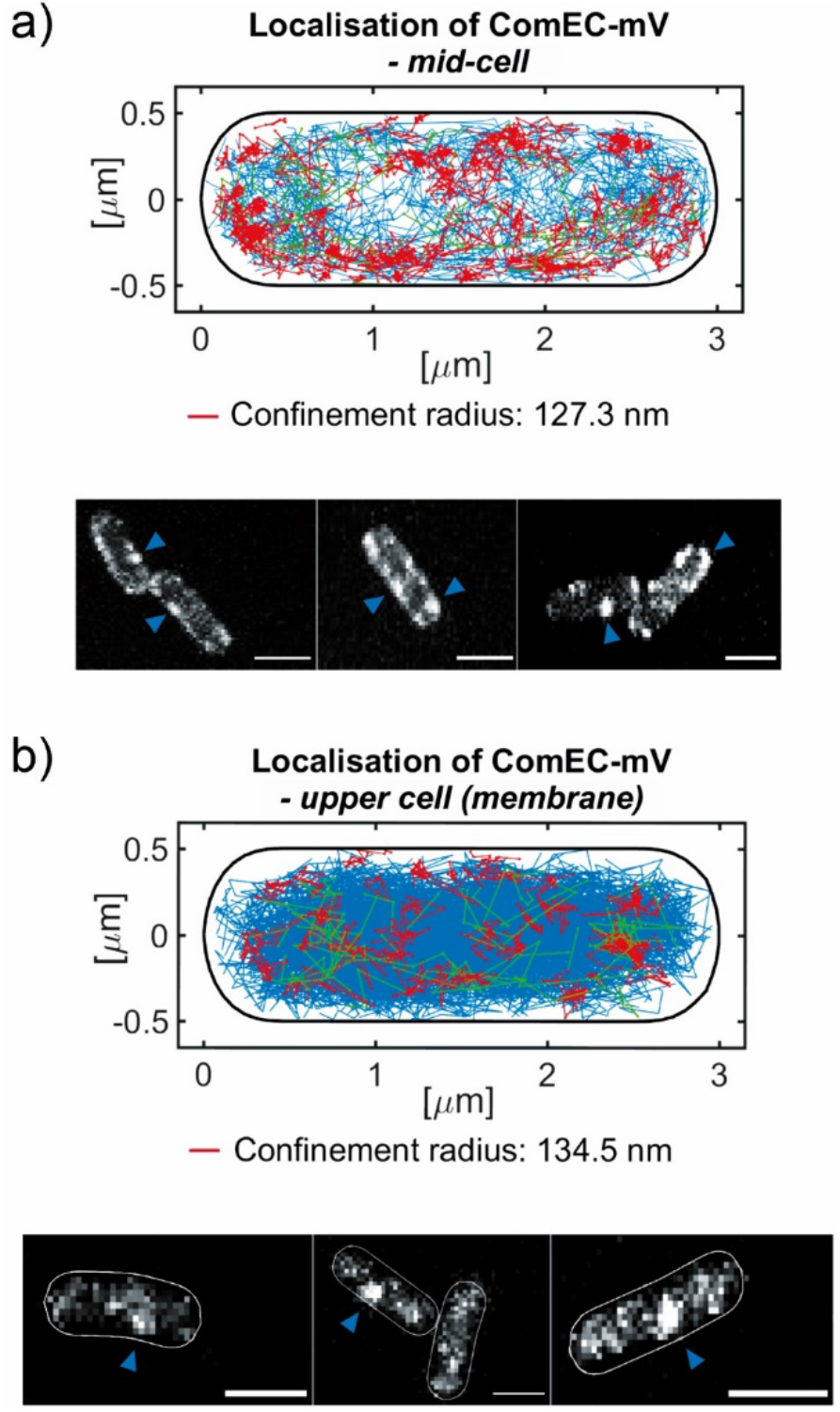
Single-molecule tracking of ComEC-mVenus. In a) the localisation of tracks is shown as an overlay in one normalised cell, molecules were tracked mid-cell (number of tracks: 548). Lower panels, exemplary t-stacks of an analysed movie with 2000 frames are shown. In b) the same experiment is shown except for the changed imaging plane (upper cell), where 1280 tracks were monitored. In red, the confined tracks are depicted, moving within a radius of either 127.3 nm (a)) or 134.5 nm (b)). The confinement radius corresponds to 3 times the standard error of the MSD (calculated from 5 intervals). Green colour indicates a change of the tracks from confined to a more mobile movement or *vice versa*; tracks are therefore considered as segmented. Steps of blue coloured tracks always exceed the confined radius; these tracks are considered as mobile. Blue arrows indicate localisation of a strong intracellular signal. Measurements were carried out in biological triplicates. White bars 2 μm.

### Diffusion of fluorescently labeled DNA can be monitored in competent *B. subtilis* cells via single-molecule tracking inside of the periplasm

We fluorescently labeled DNA according to the protocol of Boonstra *et al.* (2018) (15), and incubated *B. subtilis* cells grown to competence with a fluorescently labeled PCR-product, which displayed homology to the *B. subtilis* chromosome, flanking an erythromycin resistance gene (2300 bp, partial sequence of vector pDG1664). After an incubation time of 30 min and an additional incubation time of 40 min with DNAse, we investigated our samples by single-molecule tracking at mid-cell level. We chose strain PG876 in order to increase the number of cells entering the K-state. Initial measurements with an exposure time of 50 ms, assuming slow movement of the up to 1.5 mDa large dsDNA, resulted in mostly blurry signals but no defined point spread functions (data not shown). As a result, we recorded the signal with an exposure time of 30 ms, and to our surprise found dynamics that resemble those obtained for proteins (movie S2, movie S3). Data could be fitted best by assuming three diffusive populations, two diffusive population did not result in a sufficient quality of fitting (meaning not all measured data could be covered by a double fit). We calculated diffusion coefficients of 0.014 +/− 0.0 μm^2^/s, 0.071 +/− 0.001 μm^2^/s and 0.621 +/− 0.004 μm^2^/s for the labeled DNA (PG876, figure 8a), table 4). These data correspond to static motion seen for ComEC (table 3), to slow diffusion in case of the medium-fast population, and free diffusion of cytosolic proteins (43–45).

**Table 4:**
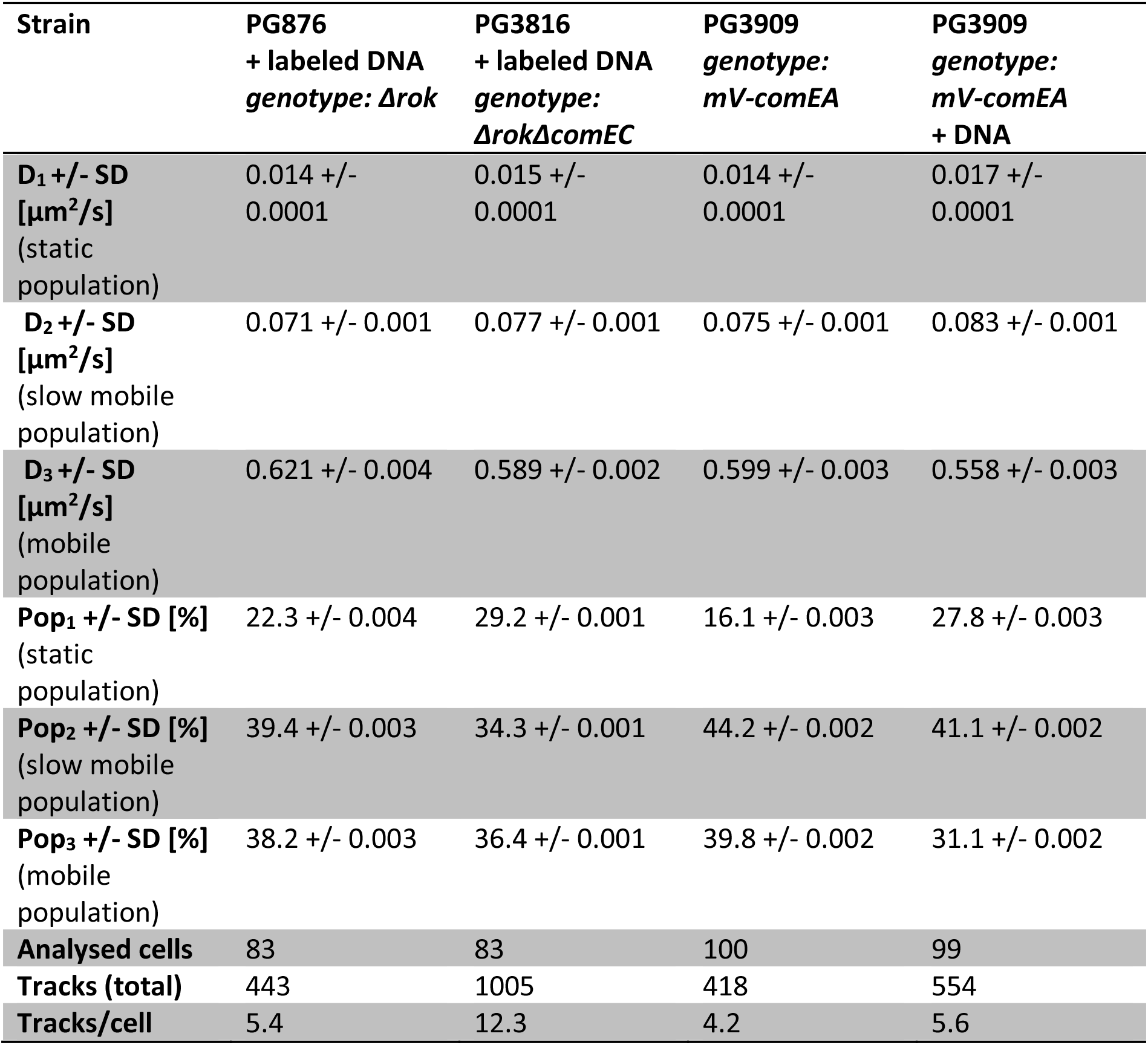
Single-molecule tracking of fluorescently labeled DNA compared to mVenus-ComEA

| Strain | PG876<br>+ labeled DNA<br>*genotype: Δrok* | PG3816<br>+ labeled DNA<br>*genotype: ΔrokΔcomEC* | PG3909<br>*genotype: mV-comEA* | PG3909<br>*genotype: mV-comEA*<br>+ DNA |
| --- | --- | --- | --- | --- |
| **D<sub>1</sub> +/- SD**<br>[μm<sup>2</sup>/s]<br>(static<br>population) | 0.014 +/-<br>0.0001 | 0.015 +/-<br>0.0001 | 0.014 +/-<br>0.0001 | 0.017 +/-<br>0.0001 |
| **D<sub>2</sub> +/- SD**<br>[μm<sup>2</sup>/s]<br>(slow mobile<br>population) | 0.071 +/- 0.001 | 0.077 +/- 0.001 | 0.075 +/- 0.001 | 0.083 +/- 0.001 |
| **D<sub>3</sub> +/- SD**<br>[μm<sup>2</sup>/s]<br>(mobile<br>population) | 0.621 +/- 0.004 | 0.589 +/- 0.002 | 0.599 +/- 0.003 | 0.558 +/- 0.003 |
| **Pop<sub>1</sub> +/- SD [%]**<br>(static<br>population) | 22.3 +/- 0.004 | 29.2 +/- 0.001 | 16.1 +/- 0.003 | 27.8 +/- 0.003 |
| **Pop<sub>2</sub> +/- SD [%]**<br>(slow mobile<br>population) | 39.4 +/- 0.003 | 34.3 +/- 0.001 | 44.2 +/- 0.002 | 41.1 +/- 0.002 |
| **Pop<sub>3</sub> +/- SD [%]**<br>(mobile<br>population) | 38.2 +/- 0.003 | 36.4 +/- 0.001 | 39.8 +/- 0.002 | 31.1 +/- 0.002 |
| **Analysed cells** | 83 | 83 | 100 | 99 |
| **Tracks (total)** | 443 | 1005 | 418 | 554 |
| **Tracks/cell** | 5.4 | 12.3 | 4.2 | 5.6 |

**Table 5:** Transformation frequencies of mutants of ComEC.

| Strain | genotype | Transformation frequency<br>+/- SD | % wild-type<br>+/- SD | % $\Delta comEC$<br>*amyE::comEC*<br>+/- SD |
| --- | --- | --- | --- | --- |
| PG001 | wild-type, PY79 | $6.407^{-005}$<br>+/- $1.425^{-005}$ | 100 | - |
| PG3722 | $\Delta comEC$ | 0 | 0 | 0 |
| PG3929 | $\Delta comEC$,<br>*amyE::comEC* | $3.345^{-005}$<br>+/- $7.329^{-006}$ | 52.2<br>+/-11.43 | 100 |
| PG3917 | $\Delta comEC$,<br>*amyE::comEC*<br>*H571A* | $4.259^{-006}$<br>+/- $2.909^{-006}$ | 6.65<br>+/- 4.5 | 12.73<br>+/- 8.7 |
| PG3918 | $\Delta comEC$,<br>*amyE::comEC*<br>*D573A* | 0 | 0 | 0 |
| PG3919 | $\Delta comEC$,<br>*amyE::comEC*<br>*D575A* | $1.055^{-005}$<br>+/- $3.413^{-006}$ | 16.5<br>+/- 5.3 | 30.57<br>+/- 9.89 |
| PG3927 | $\Delta comEC$,<br>*amyE::comEC*<br>*D668A* | $1.351^{-005}$<br>+/- $4.884^{-006}$ | 21.08<br>+/- 7.62 | 40.38<br>+/- 14.6 |
| PG3928 | $\Delta comEC$,<br>*amyE::comEC*<br>*K671A* | $2.880^{-005}$<br>+/- $4.105^{-006}$ | 44.95<br>+/- 6.4 | 86.09<br>+/- 12.27 |

After incubating the *rok* deletion strain with labeled DNA, we found a polar and mostly peripheral pattern of confined tracks (indicated in red), in addition to a fast-diffusive signal indicated in blue (PG876, *Δrok*, figure 7a)). This suggests that taken up DNA is largely present at the periphery of the cell, i.e. in the periplasm. As a control, we analysed wild-type cells (PG876) without addition of DNA, where we found an average background signal of 2 tracks/cell (see Fig. S4). Because only a low signal of 5.4 tracks/cell was detected when DNA was added, we note that many signals obtained could be due to background noise. We reasoned that an increased amount of signal might be obtained when *comEC* is deleted (in addition to *rok*), which was indeed the case (PG3816, figure 7b)). Signals changed from 445 to 1005 tracks in total (12.3 tracks/cell, see table 4). Diffusion constants for DNA in the *Δrok ΔcomEC* background were 0.015 +/− 0.0 μm^2^/s, 0.077 +/− 0.001 μm^2^/s and 0.589 +/−0.002 μm^2^/s (PG3816, figure 8b), table 4), very closely resembling those in wild type (*Δrok*) cells. Because of the much higher signal to background ratio now allows us to deduce that taken up DNA indeed shows motion within the periplasm that can be best described by three distinct mobilities.

**Figure 6:**
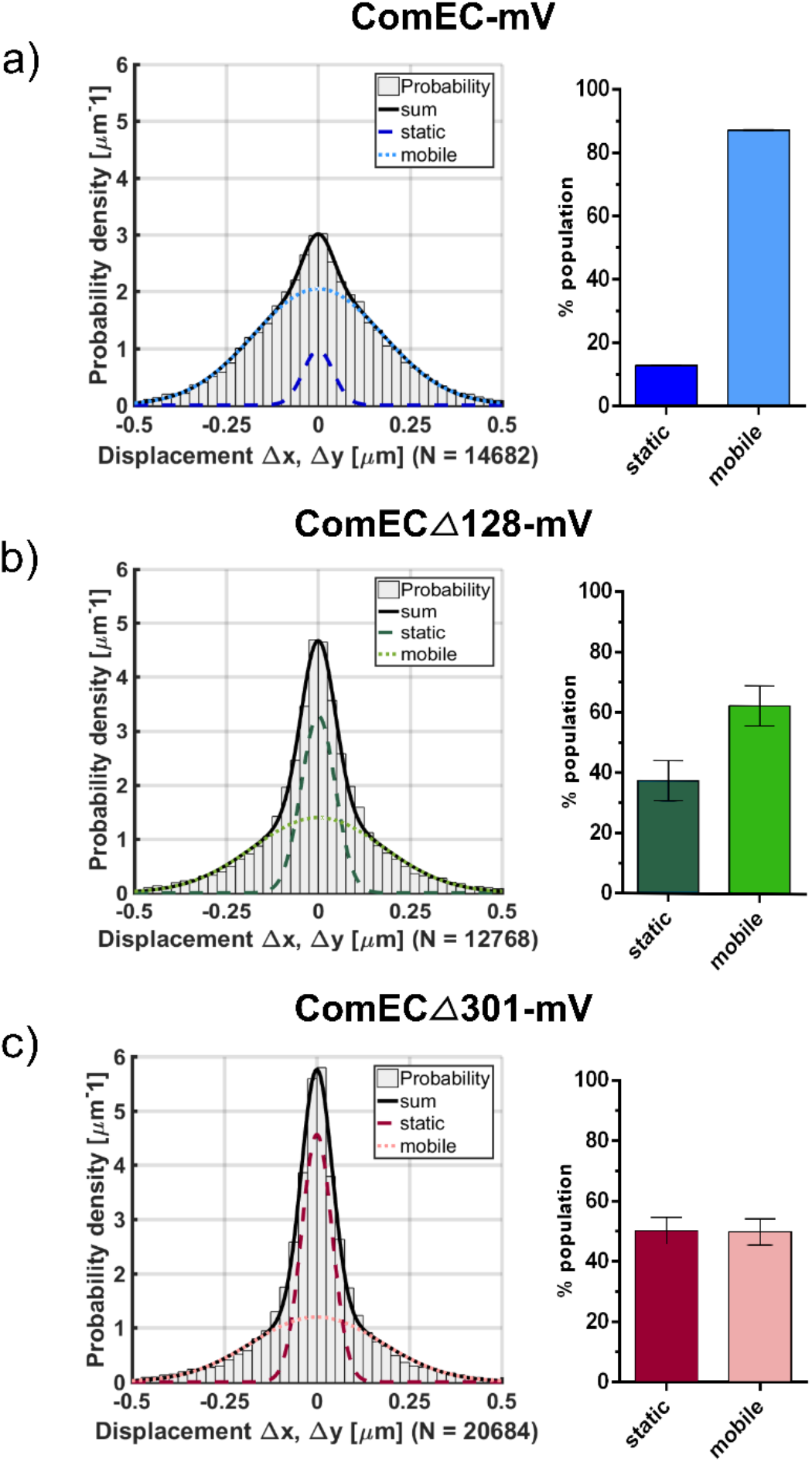
Analysis of populations of ComEC-mV and of C-terminal truncations with different mobility analysed by GMM. In a) The step-sizes of all analysed tracks for ComEC-mV (PG3817) are plotted in a histogram, according to their probability density and fitted by two Gaussian functions, whose area is then summed up to one fitting (R^2^= 1, black line). The % of static and mobile populations are depicted in a bar plot on the right. In b) the same analysis was done for ComECΔ128-mV (PG3818, R^2^= 1) and c) for ComECΔ301-mV (PG3819, R^2^= 1). Error bars of bar plots indicate standard deviation of the mean. N= number of steps.

**Figure 7:**
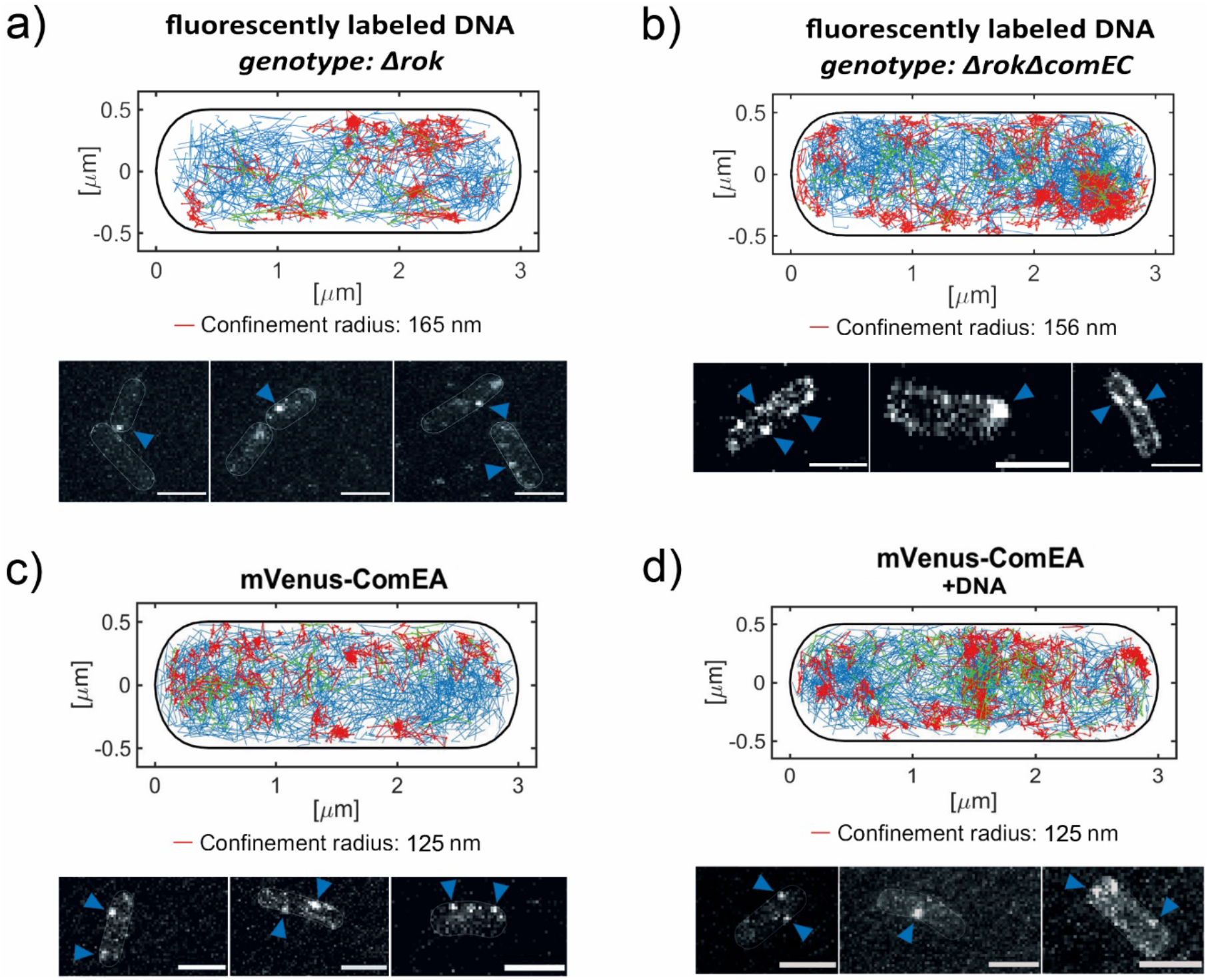
Single-molecule tracking of fluorescently labeled DNA and mVenus-ComEA. Single-molecule tracking was carried out with 30 ms exposure time. In a), the signal of *B. subtilis* PY79 cells (PG876, *Δrok*) incubated with stained PCR-product, labeled with DyLight488, is shown by plotting tracks on a standardized cell (number of tracks: 251). Lower panel exemplary t-stack of a movie, signals indicated by blue arrows. Cell meshes show the borders of the cells. b), *comEC rok* deletion strain (PG3816, n = 555) showing more distinct localisation of the signal along the membrane. c) shows localisation of mVenus-ComEA, expressed under control of the Xylose-promotor using 0.05% Xylose (strain PG3909, number of tracks: 418). In d) chromosomal DNA (20 μg/ml) was added to the mVenus-ComEA expressing cells and incubated for 1 h (n = 554). The confinement radius corresponds to 3 times the localization error. Green colour indicates a change of the tracks from confined to a more mobile movement or *vice versa*, while blue tracks consist of non-confined movements (steps) are thus of mobile molecules. Confined tracks are shown in red, moving within a radius as indicated. Cells were analysed close to mid-cell level. Scale bars 2 μm.

**Figure 8:**
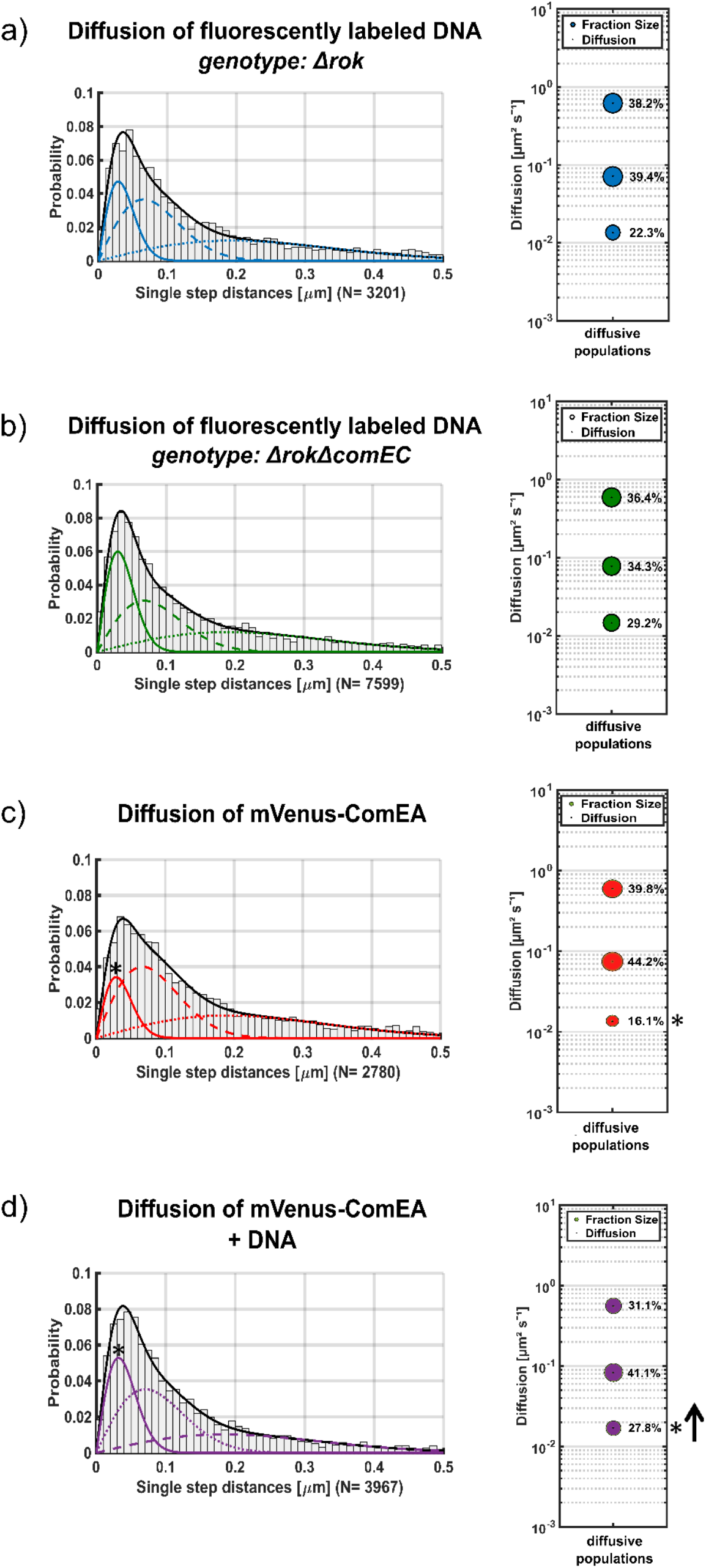
Diffusion of labeled PCR-products and of mVenus-ComEA inside competent *B. subtilis* cells. Diffusion coefficients were calculated by the squared displacement, fitting a cumulative density function to the data. Histograms in a), b), c) and d) show the probability of single step distances (Jump distance) of all acquired data, and the percentage of the diffusive populations as a bubble-plot, next to the corresponding histogram. a) and b) show single molecule tracking of labeled DNA, c) results for mVenus-ComEA. In d) chromosomal DNA was added to competent cells expressing mVenus-ComEA. Comparing c) and d), we detected an increase of the static fraction of 11.7% (fraction indicated by asterisks and black arrow in d)). For all conditions, single-molecule tracking was carried out at mid-cell level, 30 ms stream acquisition). Measurements were carried out in biological triplicates. N= number of steps.

Performing a temporal overlay of frames acquired in SMT (t-stacks of movies), we discovered a punctate localisation pattern of the labeled PCR-product along the membrane of the cells (figure 7 b), lower panel), in contrast to the fewer peripheral foci observed in wild type cells (Fig. 7a, lower panel). This observation was supported by SMT measurements, as confined tracks localised at the cell periphery (figure 7 b), upper panel, red tracks), in addition to mobile tracks, indicating a high mobility of the DNA (figure 7 b), upper panel, visualised in blue). Overall, we found a more intense polar localisation in case *comEC* was deleted (compare figure 7 a) to 7 b), lower panel), probably due to higher number of fluorescence signal in general. These findings show that in the absence of transport of DNA across the cell membrane, DNA accumulates within the periplasm, but also does so under wild type conditions, where it shows protein-like dynamics of very low to rapid diffusion.

### Movement of DNA is similar to that observed for ComEA

We wished to compare SMT data obtained for labeled DNA with the diffusion of membrane protein ComEA, which is thought to bind to DNA entering the periplasm during natural transformation of *B. subtilis*. Therefore, we created an N-terminal mVenus-fusion of the protein at the original locus (the N-terminus resides in the cytosol), which we induced with a low concentration of xylose (PG3909). Of note, even under low levels of induction, the fusion supported transformation (see figure S5). Fig. 7 shows that mVenus-ComEA was found to be mobile throughout the cell membrane. The protein accumulated at the cell-pole, where we detected a strong signal, in addition to the general localisation pattern at the membrane (see figure 7 c), movie S4). The pattern became even more obvious when mVenus-ComEA was induced with 0.5% Xylose, which we did not use for analysis as such a strong signal would distort single-molecule tracking (figure S6). Further, we incubated strain PG3909 with chromosomal DNA for 1 h, where we detected an overall stronger signal at the membrane, septum and cell pole as depicted in figure 7d). Similar to the fluorescent DNA, the diffusion of mVenus-ComEA could be best fitted using three populations. We determined diffusion coefficients of 0.014 +/− 0.0001 μm^2^/s, 0.075 +/− 0.001 μm^2^/s and 0.599 +/. 0.003 μm^2^/s (see figure 8 c, table 4), which are very similar to those determined for taken up DNA. Percentages of populations varied only little between the two conditions in which labeled DNA was investigated. About 40% of labeled DNA showed high and medium mobility, about 20% was statically positioned (see bubble plots, figure 8). In the *comEC* mutant background, more DNA molecules were static, close to 30%, while about 35% showed medium and high mobility (Fig. 8, see table 4 for exact values and standard deviations). For mVenus-ComEA, we monitored a static fraction of 16.1%, which increased (and almost doubled) to 29.2% (figure 8c and d), while diffusion coefficients did not change significantly (see table 4). The slow-mobile fraction of ComEA contained 44.2% of molecules, and only slightly decreased to 41.1% upon addition of DNA, while the fast-mobile fraction decreased from about 40 to 31% after addition of DNA. While it is not straight forward to explain the different mobilities, our results clearly show that taken up DNA can rapidly move through the *B. subtilis* periplasm, which appears to be able to act as a reservoir for DNA that can not be pumped into the cell via ComEC. Of note, we closely followed the protocol devised by the Kuipers groups to obtain comparable conditions. We cannot state if there may be differences in dynamics of DNA or of ComEA soon after addition of DNA, different from our conditions.

## Discussion

Uptake of DNA by competent bacteria has been described to occur at single cell poles, for *B. subtilis*, *Vibrio cholerae* and for *Helicobacter pylori* (14, 46–48). It has been speculated that a single entry-point into the cytoplasm may facilitate search for homology to incoming DNA on the chromosome, and for incorporation of ssDNA at corresponding loci. It has remained an intriguing question how the cell can place a multiprotein complex guiding DNA through the cell envelope into the cell at a single cell pole. In recent work, we have found evidence suggesting that the polar DNA uptake machinery in *B. subtilis* may be assembled through a diffusion/capture mechanism, from highly dynamic molecules. In our present work, we have investigated the dynamics of membrane permease ComEC and of DNA receptor ComEA, and have investigated the role of the large C-terminus of ComEC. We have also been able to follow the fate of incoming DNA into the cell via single molecule tracking.

For ComEC-mV, we found a surprisingly low number of static molecules, and 87% molecules freely diffusing within the cell membrane. Of note, we found confined motion of ComEC at many places within the cell membrane, indicating that it does not only stop for some time at the cell pole, but here, it clearly stops most often. Hahn *et al.* described the movement of fluorescent foci towards the cell pole, studying a ComGA-CFP fusion by time-lapse microscopy, postulating a putative diffusion/capture mechanism for assembly of competence proteins at the cell pole, but stated that only high-speed microscopy could reveal the true nature of the assembly (49). Using SMT, our findings support the idea of a diffusion capture model for polar localization of ComEC, in order to internalize exogenous DNA that has entered the cell via the putative pseudopilus, similar to what we have found for ATPase ComGA. (35). Intriguingly, truncating the C-terminus of ComEC strongly reduced mobility of ComEC in the membrane. It has been determined that diffusion coefficients of membrane proteins decrease with increasing numbers of TMs, but not with the size of soluble parts of membrane proteins (34). Expecting an increase in mobility of ComEC, we removed one or two membrane helices from the C-terminus (ComECΔ128-mV and ComECΔ301-mV), but found an increase of static molecules by SMT. These findings indicate that the putatively periplasmic C-terminus of ComEC is required for non-constrained diffusion in the membrane.

ComEC has only been seen to localize at the membrane in the end of exponential phase, when *B. subtilis* cells are developing competence (49) and at the cell pole in competent *Bacillus* cells, described by Kaufenstein *et al.,* (2011). Comparing the full-length fusion of ComEC-mV (PG3817) to ComECΔ128-mV (PG3818), where 128 amino acids were deleted from the C-terminus of ComEC, polar localisation decreased from about 6% to 1.6% seen by epifluorescence microscopy. Both fluorescence fusions showed a high amount of uniform (or rather diffusive) fluorescence signal in 35% and 47% of the cells (see figure 2, table 1). In contrast, using SMT, we were able to detect a clear membrane-associated localization pattern of ComEC-mV and ComECΔ128-mV (figure 5a, figure S2). In case of ComECΔ301-mV, a large part of 301 amino acids was truncated, leading to a signal similar to a membrane-stain (figure 1), showing a distinct punctate localization pattern along the membrane. However, we detected a large, about three fold increase of the slow mobile/static fraction, showing that is spite of a loss of polar clustering, the truncation greatly lost diffusive mobility within the entire cell membrane.

To obtain further knowledge on the function of the C-terminus, we mutated several conserved amino acids of ComEC, identified by Baker *et al.* (2016), in order to study the putative exonuclease function possibly carried out by its C-terminus (figure 4). We found that an aspartate residue, **D**573, part of the putative zinc-binding motif HA**D**QDH, is essential for transformation. This finding supports the idea of ComEC being the putative exonuclease of the system. Unfortunately, we have been unable to provide *in vitro* assays to finally prove the theory, hampered by DNA contamination and insolubility when we expressed putative soluble parts of ComEC in *E. coli* (unpublished data).

Most importantly in this work, we have been able to follow the dynamics of fluorescently labeled DNA in real time, for which we have not found a precedence in the literature. We determined that a 2300 bp fragment of DNA (MW= 1.5 mDa) shows three distinct patterns of movement: we observed a slow mobile/static fraction, which may represent DNA that is taken up by the pseudopilus, possibly a slow event. This fraction mildly increased in cells lacking ComEC, indicating that ComEC affects the dynamics of static DNA molecules to only a small degree. We found two additional populations, one having an intermediate (low) diffusion constant, and one having a diffusion constant that is comparable to that of freely diffusing cytosolic as well as membrane proteins (50, 51). In order to better understand the nature of the three populations, we followed the motion of DNA receptor ComEA by single molecule tracking. We found that also ComEA moves as three distinct populations and curiously, all three having very similar diffusion coefficients as those determined for DNA (see figure 8, table 4). ComEA showed confined motion at many places within the cell membrane, in addition to polar accumulation like ComEC or ComGA. The three populations of ComEA could be described as mobile molecules diffusing freely without bound DNA, molecules that move together with DNA, and molecules that bind to DNA that is moved through the cell wall by the pseudopilus. This idea is supported by the finding that the fast mobile fraction of ComEA moves with an average diffusion constant similar to that of single membrane proteins (34), while that of the medium fast and slow mobile fractions is much lower. Likewise, 1.5 mDa linear DNA might move like a large cytosolic protein through the periplasm by itself. On the other hand, it is striking that diffusion constants between the three DNA fractions and those of ComEA have such similar values. Of note, when large DNA strands are added to competent *B. subtilis* cells, the DNA uptake machinery generates on average 25.000 bp DNA fragments (13, 52, 53) through the activity of NucA endonuclease (54). Thus, low mobility DNA could be ComEA bound to 2300 bp DNA, and the fast population shorter DNA fragments bound to ComEA. However, addition of DNA to competent *B. subtilis* cells did not lead to strong changes in ComEA dynamics, except for an increase in the static fraction of ComEA. Note that a portion of *B. subtilis* cells entering stationary phase actively secrete chromosomal DNA, and fragments thereof (55), such that ComEA is likely also engaged in some DNA binding in the absence of externally added DNA. Thus, alternatively, the two mobile fractions could be due to few or many ComEA molecules bound to a single DNA fragment, because ComEA can bind cooperatively to incoming DNA (48).

We interpret these data in a way that taken up DNA is directly bound by ComEA (Fig. 9). Uptake of DNA through the pseudopilus may be a slow event, leading to statically positioned ComEA and DNA. Only a small fraction of DNA is directly taken up further through ComEC, but DNA can be buffered and stored by ComEA (Fig. 9). The protein is very mobile during competence, similar to ComEA of *Vibrio cholerae*, despite the fact that *B. subtilis* ComEA is likely an integral membrane protein, and not a periplasmic receptor as it has been found for *V. cholerae* (21, 47). Because of similar diffusion constants, we favour the view that all taken up DNA is bound to ComEA, some freely diffusing, some being in contact with the uptake machinery, e.g. ComEC, or getting targeted to the pole by other factors.

**Figure 9:**
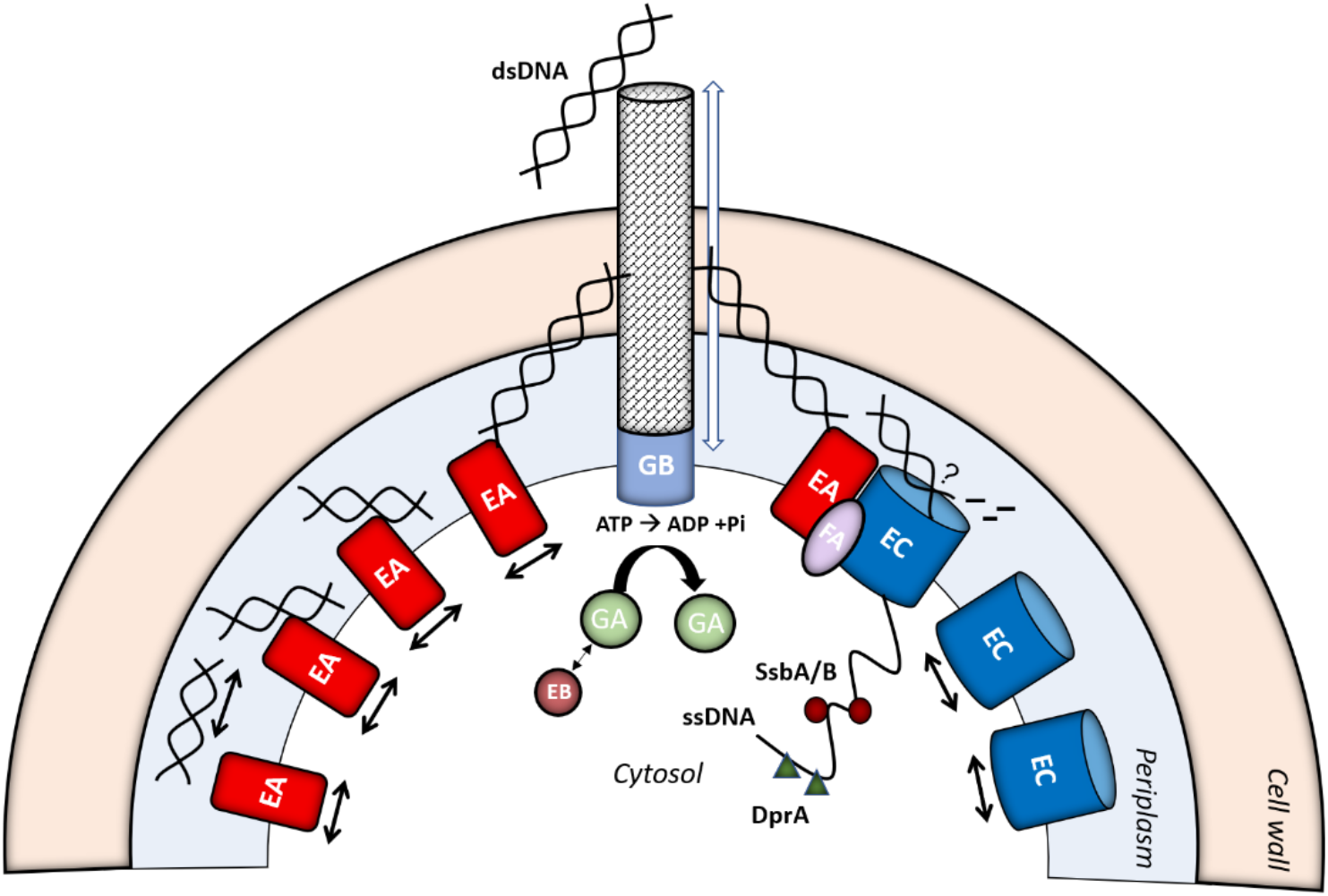
Model for DNA-uptake of *B. subtilis.* Double-stranded DNA (dsDNA) is transferred into the periplasm via rapid assembly and disassembly of the putative pseudopilus at the cell pole (indicated by weight arrow), whose anchor might be the membrane protein ComGB. ComGA is guided to the cell pole by ComEB, where it provides the energy for pilus assembly/disassembly. ComEA binds the dsDNA in the periplasm, where it can either hand over the DNA to ComEC, and ssDNA is transferred to the Cytosol with energy provided by ComFA (complex on the right), or diffuses along the membrane to act as a kind of reservoir for transforming DNA (loaded ComEA on the left). ComEC and ComEA are diffusing freely until a complex is formed at the cell pole when DNA is present (diffusion indicated by black arrows). Probably, free DNA is present in the periplasm in addition, in case it has not been bound by ComEA yet. Once ssDNA enters the Cytosol, it is coated by single-strand DNA binding proteins SsbA, SsbB and DprA, followed by integration into the chromosome.

Our findings show that unlike *V. cholerae*, ComEA is not accumulating only at the competence pole upon addition of DNA to receive incoming DNA, but remains diffusing through the membrane. The fact that diffusion constants and fraction sizes of DNA do not change markedly between wild type and *comEC* mutant cells suggests that ComEA can bind to excess incoming DNA that cannot be directly transported into the cytosol, and acts as a buffer or storage system, allowing DNA to enter at later time points. A similar function has been reported for ComE of the gram-negative *Neisseria gonorrhoeae*, where the homologue of ComEA (ComE) is considered to bind DNA in the periplasm and act as a reservoir for taken-up DNA during natural transformation (56, 57). Clearly, most of the tDNA accumulates in the periplasm, and only few molecules are transported into the cytosol, or otherwise the difference between wild type and *comEC* mutant cells would have been much bigger. We assume, that the signal we detected must have been present in the periplasm, as cells were treated with DNAse and washed, removing unspecific DNA from the cell surface. It is not clear whether the signal we detected in the wild type (PG867), was localised in the cytosol in addition, as bleed-through from the lower cell level could have also caused the detection of tracks localised at the cell center (see figure 7a)).

Our findings further improve our understanding of the *B. subtilis* competence machinery, a highly dynamic assembly of proteins, likely set up predominantly by a rapid diffusion/capture mechanism. Even large DNA molecules show high mobility within the bacterial periplasm, ensuring that it can also efficiently find the polar ComEC channel for eventual uptake into the cytosol. The idea of ComEA buffering DNA uptake agrees with findings from V. cholerae where DNA transport through the outer membrane and through the cell membrane has been shown to be spatially but not temporally coupled (48).

## Acknowledgements

We would like to thank Lisa Stuckenschneider for providing vector pHJDS-mV, Luis M. Oviedo Bocanegra for supporting our Single-molecule tracking analysis, and Daniel J. Rigden for providing the model of the C-terminus of *B. subtilis* ComEC. In addition, we thank David Dubnau for providing deletion strains of competence proteins. This work was supported through the state of Hessen (central funds for University of Marburg) and the Deutsche Forschungsgemeinschaft (DFG).

